# Remdesivir-ivermectin combination displays synergistic interaction with improved *in vitro* antiviral activity against SARS-CoV-2

**DOI:** 10.1101/2020.12.23.424232

**Authors:** Laura N Jeffreys, Shaun H Pennington, Jack Duggan, Claire H Caygill, Rose C Lopeman, Alastair F Breen, Jessica B Jinks, Alison Ardrey, Samantha Donnellan, Edward I Patterson, Grant L Hughes, David W Hong, Paul M O’Neill, Ghaith Aljayyoussi, Andrew Owen, Stephen A Ward, Giancarlo A Biagini

## Abstract

A key element for the prevention and management of COVID-19 is the development of effective therapeutics. Drug combination strategies of repurposed drugs offer several advantages over monotherapies, including the potential to achieve greater efficacy, the potential to increase the therapeutic index of drugs and the potential to reduce the emergence of drug resistance. Here, we report on the *in vitro* synergistic interaction between two FDA approved drugs, remdesivir and ivermectin resulting in enhanced antiviral activity against SARS-CoV-2. These findings warrant further investigations into the clinical potential of this combination, together with studies to define the underlying mechanism.

## INTRODUCTION

At the time of writing, the World Health Organisation (WHO) has reported more than 211 million cases of COVID-19, and more than 4.4 million deaths (1). There remains a clear need for therapeutic strategies with activity against SARS-CoV-2. Potential therapeutic strategies may include the repurposing of existing drugs as well as the discovery of novel therapies. Thousands of clinical trials are currently underway, with therapeutic approaches involving direct-acting antivirals, for the prevention of virus replication, and host-directed therapies aimed at mitigating against the disease pathology (2, 3).

Combination therapies can offer several advantages over monotherapies. They have the potential to achieve greater efficacy, to increase the therapeutic index of drugs and to reduce the emergence of drug resistance. Strategies to identify effective combination therapies are emerging, with several laboratories reporting *in vitro* combination screens (4) and *in vivo* animal combinations studies (5). In a recent clinical trial, baricitinib administered in combination with remdesivir was found to be superior, and to elicit fewer adverse effects, compared to either drug in isolation (6). Importantly, even in the absence of synergistic activity, an additive interaction between two drugs with separate mechanisms of action may profoundly reduce the speed at which drug resistance is established.

Both remdesivir and ivermectin have received attention for the treatment of COVID-19. Remdesivir is a prodrug C-adenosine nucleoside analogue that inhibits the viral RNA-dependent, RNA polymerase. Remdesivir was shown early in the pandemic to display *in vitro* antiviral efficacy against SARS-CoV-2 (7). In a double-blind, randomized, placebo-controlled trial, intravenous administration of remdesivir showed superiority relative to placebo in shortening the time to recovery in adults who were hospitalized with COVID-19 (8). However, other studies have suggested that its impact may be negligible (9), and on 20^th^ November 2020 the WHO issued a conditional recommendation against the use of remdesivir in hospitalised patients (irrespective of disease severity) because there is no evidence supporting an improvement in survival or other outcomes in these patients.

Ivermectin is an anti-parasitic which is active against a wide range of parasites, including gastrointestinal roundworms, lungworms, mites, lice, hornflies and ticks (10). Ivermectin is reported to exhibit broad spectrum anti-viral activity against a wide range of RNA and DNA viruses (11). Recently, ivermectin was also shown to display anti-viral activity against SARS-CoV-2 (12), but approved doses are not expected to be high enough to achieve *in vitro*-defined target exposures systemically (13). Several clinical trials are now evaluating the potential of ivermectin for both prophylaxis and treatment of COVID-19, but the low exposures make the anti-inflammatory and/or immunomodulatory mechanisms of action more plausible than a direct antiviral activity of the monotherapy (14). In particular since studies with SARS-CoV-2 in Syrian Golden Hamsters showed an impact upon disease pathology in the absence of any effect on viral titres (15).

Here, we present the Fractional Inhibitory Concentration Index (FICI), and report a synergistic interaction between remdesivir and ivermectin resulting in improved *in vitro* antiviral activity against SARS-CoV-2. The data are discussed in the context of current therapeutic efforts against COVID-19.

## MATERIALS AND METHODS

### SARS-CoV-2 Strain

SARS-CoV-2/Human/Liverpool/REMRQ0001/2020 was isolated from a nasopharyngeal swab from a patient in Liverpool and passaged a further 4 times in Vero E6 cells. The mapped RNA sequence has previously been submitted to Genbank, accession number MW041156.

### Vero E6 cell culture and plate preparation

Vero E6 cells were maintained in complete EMEM (EMEM supplemented with 10% heat-inactivated (HI) FBS [Gibco; 10500-064] and 1% penicillin/streptomycin [Gibco; 15140-122]) in T150 flasks (Thermo Fisher Scientific) at 37°C with 5% CO_2_. Cells were seeded in resting EMEM at 1 × 10^5^ cells/well in 96-well plates (Grenier Bio-one; 655090). Plates were then incubated for 20 hours at 37°C with 5% CO_2_ to allow the cells to reach 100% confluence. The resting minimal medium was then removed, and the cells used for downstream applications.

### Concentration-response for remdesivir and ivermectin against SARS-Cov-2

VERO E6 cells were treated in triplicate with either drug in minimal medium at 25.00 μM, 8.33 μM, 2.78 μM, 0.93 μM, 0.31 μM, 0.10 μM and 0.03 μM (DMSO maintained at 0.25%) or control media, as appropriate. The plates were then incubated at 37°C with 5% CO_2_ for 2 hours. The minimal media containing the experimental compounds and the control media was then removed. 50 μL minimal media containing SARS-CoV-2 (MOI of 0.005), 100 μL 2× semi-solid media and then 50 μL minimal media containing experimental compounds and control media was added to each well, as appropriate. After 48 hours, 4% v/v paraformaldehyde was added to each well and the plate incubated for 1 hour at room temperature. The medium was removed and cells were stained with crystal violet. Cells were washed three times with water and cytopathic viral activity was determined by measuring absorbance of each well at 590 nm using a Varioskan LUX microplate reader (Thermo Fisher Scientific). Z′ was calculated for each plate using the uninfected/untreated controls and infected/untreated according to equation 1.

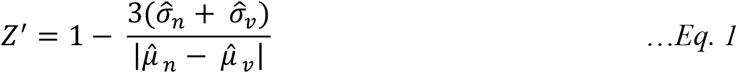

Where 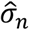 and 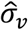 represent the standard deviation of the non-viral and viral controls respectively, while 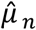 and 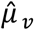 represent the corresponding means of these controls. Drug activity was expressed as a percentage of inhibition of viral growth relative to the uninfected/untreated control (100% inhibition of viral cytopathic activity) and the infected/untreated control (0% inhibition of viral cytopathic activity) on that plate.

### Fractional Inhibitory Concentration Index for remdesivir-ivermectin combinations against SARS-CoV-2

Following the assessment of the inhibitory effect (EC_50_) of remdesivir and ivermectin monotherapy on the cytopathic viral activity of SARS-CoV-2, the FICI was determined using the method developed by Berenbaum (16), using data from four independent biological replicates. Drug stocks were created in DMSO to provide a stock sufficient to produce a top concentration of 25 μM for each biological replicate. Drugs were then combined to generate mixed ratios of 1:0, 0.8:0.2, 0.6:0.4, 0.4:0.6, 0.2:0.8 and 0:1.0. Fixed ratios were then diluted across an 8-point concentration range 1:2 (DMSO maintained at 1%) to generate concentration-response data for each ratio. Z′ was calculated and quality control implemented as above.

Ratio dilutions were performed in a single 2mL deep-well plate, and added in parallel to three 96-well plates for each biological replicate. One additional plate which was not inoculated with virus was included to observe drug toxicity. Compound incubation and viral addition was performed as described above. Interpretation of FICI (FICI<=0.5 = synergy; FICI>4.0 = antagonism; FICI>0.5-4 = no interaction) was based on guidance provided by the *Journal of Antimicrobial Chemotherapy* (17).

## RESULTS

Initial experiments were conducted to determine the anti-SARS-CoV-2 activity of ivermectin and remdesivir in isolation. For each compound a robust 4-parameter fit was generated (Figure 1). For ivermectin the EC_50_ was 2.4 ± 1.1 μM and for remdesivir the EC_50_ was 1.3 ± 2.1 μM (geometric mean ± geometric standard deviation, *n*= 3).

**Figure 1.**
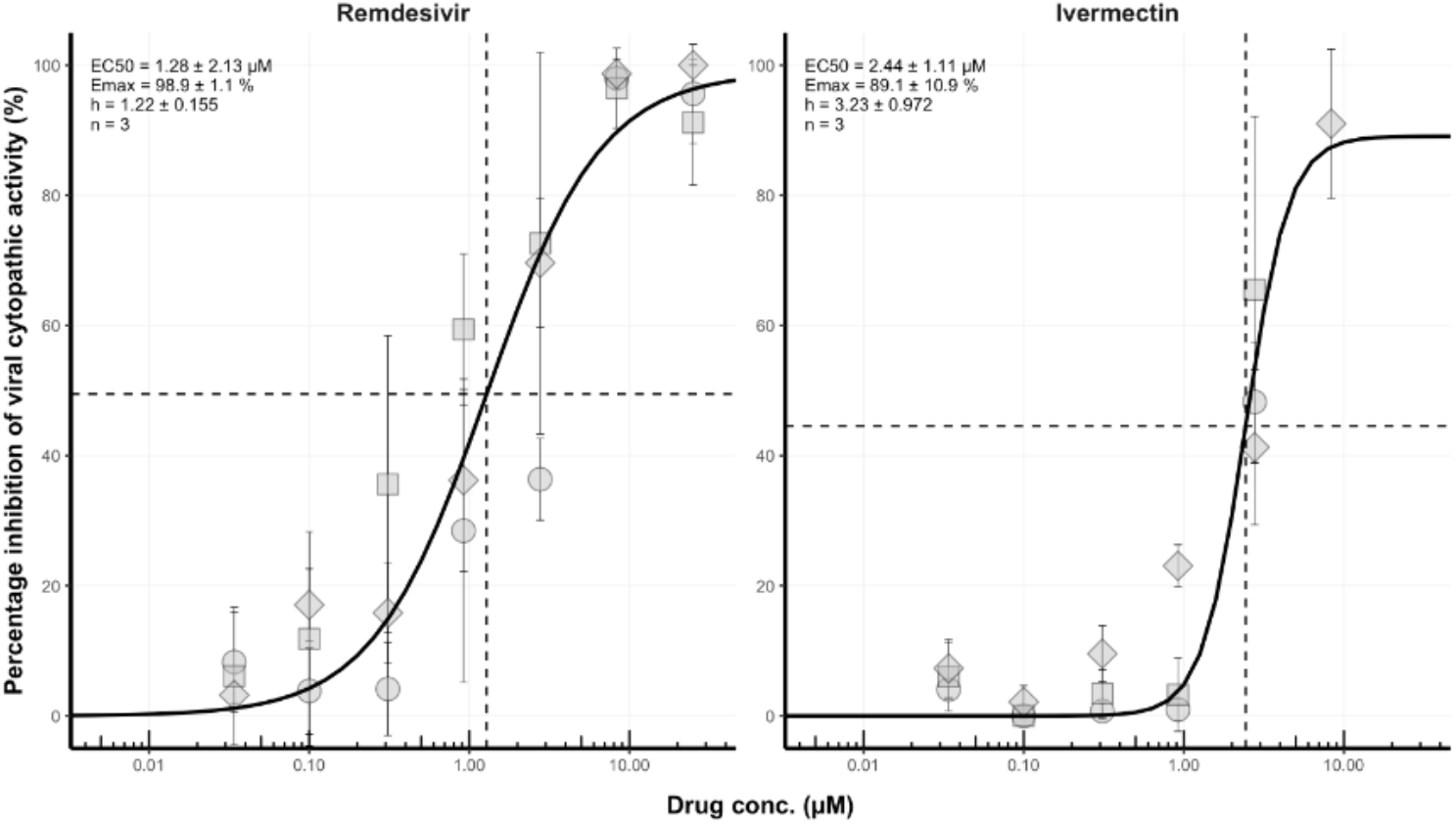
Concentration-effect relationship for the inhibition (%) of SARS-CoV-2 cytopathic activity for remdesivir and ivermectin. For each compound, activity was expressed relative to uninfected/untreated controls (100% inhibition of viral cytopathic activity) and infected/untreated controls (0% inhibition of viral activity). For each compound, we assessed activity at 25.00 μM, 8.33 μM, 2.78 μM, 0.93 μM, 0.31 μM, 0.10 μM and 0.03 μM in triplicate. Data points impacted by drug toxicity were removed automatically. Non-linear regression using an E_max_ model was performed on data taken from three independent biological replicates in order to generate concentration-effect predictions (solid black lines). For each compound, EC_50_ values, hillslope and replicate number (n) are shown. Dashed lines represent the EC_50_ of each compound. Circles, triangles and diamonds represent individual biological replicates and error bars represent standard deviation calculated from technical triplicates.

We subsequently determined the combination interaction between remdesivir and ivermectin by isobologram. The 0.2:0.8 (remdesivir:ivermectin [5 μM:20 μM]) ratio and 0.4:0.6 ratio (10 μM:15 μM), demonstrated synergy (FICI<0.5) across all 4 biological replicates (Figure 2). 3/4 biological replicates demonstrated synergy for the 0.6:0.4 ratio, and 1/4 biological replicates demonstrated synergy for the 0.8:0.2 ratio (Figure 2). For all biological replicates which did not meet the defined threshold hold of synergy, their activity did consistently exceed the predicted effect assuming a purely additive relationship (Figure 2).

**Figure 2.**
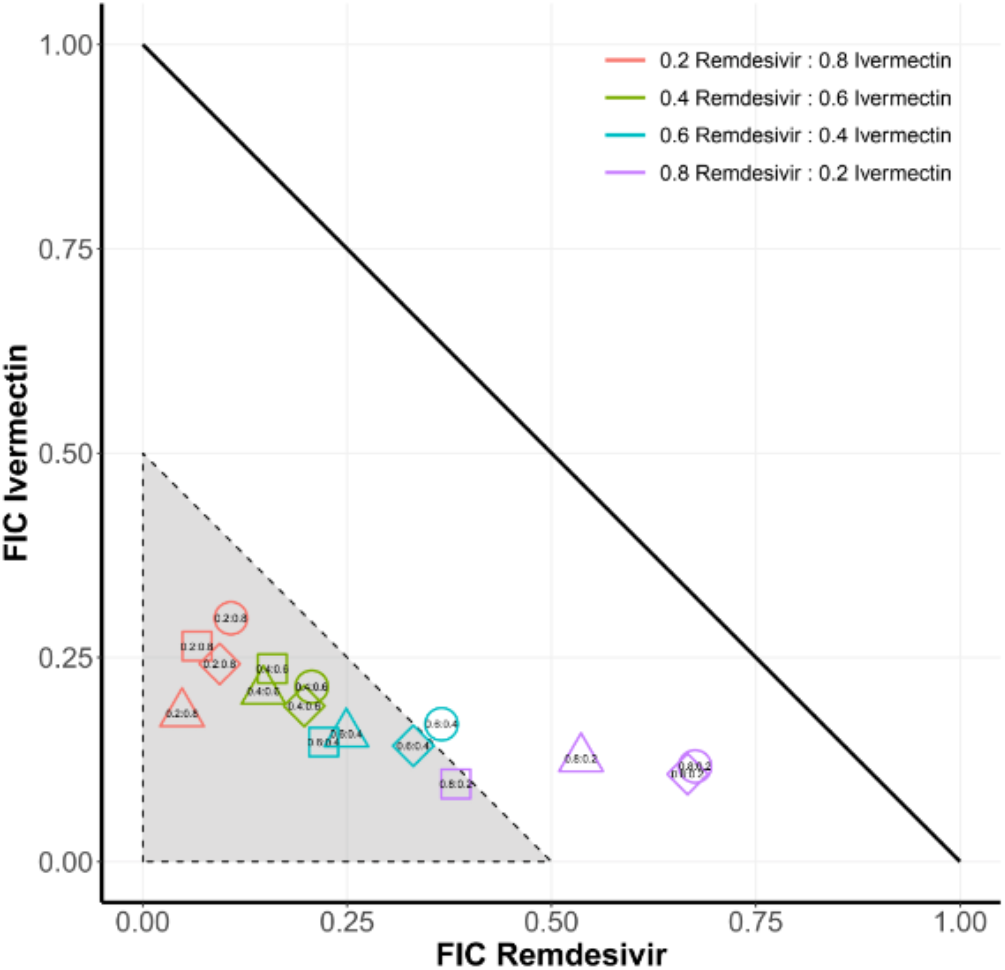
Ivermectin and remdesivir display synergistic interaction. Using EC_50_ values ranges of ivermectin and remdesivir were analysed for synergy from 8-point sigmoidal curves. Data are presented for fixed concentrations at 25 μM (corresponding to 1.0), 20 μM (0.8), 15 μM (0.6), 10 μM (0.4) and 5 μM (0.2). The area indicating synergy (FIC <=0.5) is shown in grey. Squares, triangles, diamonds and circles correspond to individual biological replicates, each derived from technical triplicates.

## DISCUSSION

Here, we described the synergistic interaction between two FDA approved drugs resulting in enhanced *in vitro* antiviral activity against SARS-CoV-2. Although combination therapy offers a number of advantages to monotherapy, genuine descriptions of synergy are relatively infrequent (18). Despite thousands of combination experiments having been performed, there have been very few reports of validated synergistic interactions against SARS-CoV-2 (4, 19). At this stage, the mechanism underpinning the synergistic interaction between remdesivir and ivermectin is unclear (7, 12).

Based on available pharmacokinetic data, concentrations required to inhibit viral replication will not be achieved in human plasma (13). However, modelling does indicate that inhibitory concentrations may be achieved in lung (13), although this has been challenged (20). Nonetheless, a combination containing ivermectin may offer the opportunity to exploit the anti-inflammatory and/or immunomodulatory activity of this agent, while simultaneously augmenting the antiviral activity of RNA polymerase inhibitors. If this synergy is also evident with orally bioavailable polymerase inhibitors such as favipiravir or molnupiravir, the approach may be more widely exploitable for community deployment.

Data presented here suggest that remdesivir in combination with ivermectin may enhance antiviral activity, reduce the risk of side effects and reduce the cost of treatment. Further investigation is now required to determine whether the observed synergistic interaction can be replicated in animal models, and if these results are favourable, whether the finding can be translated to the clinic. It remains to be determined whether the observed synergistic remdesivir-ivermectin interaction will result in a favourable shift in the plasma C_max_/EC_90_ ratio. Nevertheless, the underpinning mechanisms for this synergy warrant further investigation so that this pharmacodynamic phenomenon can be exploited for the development of optimal drug combinations.

## TRANSPARENCY DECLARATIONS

A.O. is a Director of Tandem Nano Ltd. A.O. has received research funding from ViiV, Merck, Janssen and consultancy from Gilead. These associations had no influence over the content of the current manuscript. P.O.N. is currently engaged in a collaboration with Romark LLC but this interaction had no influence over the content of the current manuscript. No other conflicts are declared by the authors.

